# Hand Gesture Prediction via Transient-phase sEMG using Transfer Learning of Dilated Efficient CapsNet: Towards Generalization for Neurorobotics

**DOI:** 10.1101/2022.02.25.482002

**Authors:** Eion Tyacke, Shreyas P. J. Reddy, Natalie Feng, Rama Edlabadkar, Shucong Zhou, Jay Patel, Qin Hu, S. Farokh Atashzar

## Abstract

There has been an accelerated surge to utilize the deep neural network for decoding central and peripheral activations of the human’s nervous system to boost up the spatiotemporal resolution of neural interfaces used in neurorobotics. Such algorithmic solutions are motivated for use in human-centered robotic systems, such as neurorehabilitation, prosthetics, and exoskeletons. These methods are proved to achieve higher accuracy on individual data when compared with the conventional machine learning methods but are also challenged by their assumption of having access to massive training samples.

**Objective:** In this letter, we propose Dilated Efficient CapsNet to improve the predictive performance when the available individual data is very minimum and not enough to train an individualized network for controlling a personalized robotic system.

**Method:** We proposed the concept of transfer learning using a new design of the dilated efficient capsular neural network to relax the need of having access to massive individual data and utilize the field knowledge which can be learned from a group of participants. In addition, instead of using complete sEMG signals, we only use the transient phase, reducing the volume of training samples to 20% of the original and maximizing the agility.

**Results:** In experiments, we validate our model performance with various amounts of injected personalized training data (25%-100% of transient phase) that is segmented once by time and once by repetition. The results of this paper support the use of transfer learning using a dilated capsular neural network and show that with the use of such a model, the knowledge domain learned on a small number of subjects can be utilized to minimize the need for new data of new subjects while focusing only on the transient phase of contraction (which is a challenging neural interfacing problem).

## I. Introduction

In the U.S., there were nearly 2 million people living with limb loss [1], and by estimation, the total number of amputees will approximately exceed 4 million within 30 years [2]. Among all amputations, upper limb loss has less frequent occurrence but is reported a higher rejection rate on its commercial prosthesis [3],[4]. The deviation between amputees’ perception and prosthetic accuracy, as well as the poor functionality and discomfort of the upper limb prosthesis both lead to the high rejection rate [5], calling for a more intelligent upper-limb prosthesis system with more accurate detection of amputees’ intended movements and more agile response to amputees’ muscle-activity signal.

In the literature, surface electromyography(sEMG) signals [6]–[12] have been extensively utilized to realize the prosthetic control in a non-invasive manner. Hand gesture classification through the sEMG signals is thus considered a prominent approach. Conventionally, researchers have been focused on extracting spectral and temporal features from EMG signals and feeding them into classical machine learning algorithms [7]–[10] such as Linear Discriminant Analysis (LDA), Support Vector Machines (SVM), and Random Forest (RF). However, those algorithms require manual feature extraction and can only achieve relatively low accuracy with a large number of gestures to be classified.

Recently, deep neural networks [11]–[15] have been introduced to improve the performance of sEMG-based hand gesture recognition tasks specifically on individualizd modeling which means that a new model would be needed for any new subject and possibly for any new use of the system by any new subject. This challenges the practicality and adoption by the end-user, and is the main challenge targeted in this paper. Among neural network systems, Convolutional Neural Networks (CNNs) [11],[12] help eliminate the need for manual feature extraction and improve the model performance. Recurrent Neural Networks (RNN) [13],[14] perform even better due to their ability to capture temporal dynamics. The hybrid models that incorporate CNN and RNN are also applied to leverage the advantages of both techniques [15]. Those deep-learning methods help achieve higher classification accuracy on a larger number of gestures but also require a much larger amount of data fed as training samples.

Unfortunately, a large volume of sEMG signals is hard to collect in a real-life situation, let alone the fact that sEMG signals have inherent variability characteristics [16]. By nature, the neural drive to muscles is time-dependent and stochastic, and the neural control strategies between different users and amputation conditions can also cause a certain degree of variability. Consequently, the models trained on specific subjects may not be consistently reused over time, and the model that works for one user will not perform well for another user. In gesture detection, there comes the convention of establishing user-specific models. While in a real-life situation, a single user of prosthetic products is usually unable to provide sufficient data that deep neural networks need. Frequent retraining and recalibration of prosthesis models [17] is one way to compensate for the shortage, while the inconvenience it brings also prompts the high rejection rate in the upper limb rehabilitation system.

To augment the performance of such systems there are hardware and software innovations. For examples, using a dens matrix of electrodes and recurrent neural network, we have recently shown that the performance of individualized model can be significantly improved. In this current paper we focus on using conventional bipolar EMG (not high-density) and we propose a novel transfer learning algorithm that can cognitively record the knowledge domain from a group of subject (regarding the neurophysiological maping of muscle activation to motion) and extrapolate that knowledge for new users significantly reduicing the need for having access to new data for individual uses. Transfer learning, which serves as a method to deal with data deficiency while ensuring classification accuracy, was recently introduced into sEMG hand gesture detection [18]–[20]. It stores the knowledge gained from pre-trained models and transfers the knowledge (learned parameters) to the current model. In this way, it vastly reduces the training samples needed and, at the same time, improves the model performance by common knowledge gained in advance. Currently, most studies conducted on sEMG transfer learning are based on convolutional neural networks [19],[20]. Though it reduces the need for retraining and recalibration, the pre-trained model still needs large amounts of data to achieve good performance.

In this letter, as the first step in this work we utilize only transient phase signals for hand gesture detection, which further reduces the data volume required for training and allows the current hand gesture “classification” problem becoming “prediction.” From an engineering perspective, a voluntary muscle contraction can be divided into the transient phase, the steady-state, and the descending phase. The transient phase corresponds to the bursts of myoelectric activity triggered by sudden muscular effort; the steady-state is associated with the myoelectric signal produced by stable muscle contractions; the descending phase relates to the muscular relaxation after a gesture is achieved. In other words, the transient phase is the signal recorded after initiation of the contraction and before the subject holds the same movement in place for the plateu data collection. In previous research [6]–[20], most of the existing state-of-the-art models are trained and test on the complete sEMG signals or only use steady-state signals. This would be a simpler decoding problem to solve due to the reduction of stochasticity in the signal which would eventually help to improve model performance. However, transient phase signals are often ignored due to their unstable appearance. However, in contrast to the under-utilization situation, transient phase signals have actually been observed to possess a deterministic structure [21], suggesting orderly recruitment of motor units and the potential of including descriptive information of intended movements [22]. By utilizing the transient phase, we can reduce training signals to at least 20% of the original amount, and predict the hand gesture before movements are performed.

Nevertheless, achieving high accuracy with deep learning methods usually requires large amounts of training samples. To maintain and even enhance the predictive accuracy with much less training data, we apply transfer learning on the Dilated Efficient CapsNet model. To summarize: 1) we use Capsule Neural Network instead of conventional CNN to save inputs as vectors and thus, detect wherever the hand gesture signal is within the sEMG data; 2) we use Efficient CapsNet instead of CapsNet, removing the decoding block which is unnecessary for preprocessed signal data, and thus, reduces a huge number of training parameters and simplifies the model complexity; 3) We use dilation in each convolutional layer of the convolutional block to include edge information into prediction and thus increase the predictive accuracy by about 2% each subject. To evaluate the robustness and superiority of the proposed model, we also conduct a comparative study with other conventional deep neural networks using different percentages of the transient-phase signals as training data. The main contributions of this work are as follows:

### Contribution 1

This letter introduces Dilated Capsule Convolutional Networks to sEMG-based hand gesture prediction for the first time, evaluating the performance of the model for various injection of the new data. By saving vectors instead of scalars, CapsNet enlarges the model’s ability to detect unique underlying physiological information associated with each gesture.

### Contribution 2

By using different percentages of transient phase signals as training data, we reduce the amount of training sample to 5% ∼ 20% of the original need, and significantly enhance the agility and temporal resolution of prosthesis systems.

### Contribution 3

By proposing transfer learning techniques of Dilated Efficient CapsNet, we provide a model architecture that minimizes the amount of training data and can achieve about 70% accuracy when predicting gestures only based on 5% of the original training data.

## II. Data

### A. Data Acquisition

Ninapro [23] is an open-source project that aims to help EMG prosthetics research through the publicly available sEMG dataset. In this work, we leverage its second subdatabase (DB2), and use the section of “Exercise set B”. This dataset includes the sEMG signal [24] of 40 subjects (all intact; 28 males, 12 females; 34 right-handed, 6 left-handed; age 29.9 ± 3.9 years) performing 17 hand gestures shown in the Fig. 1 (8 isometric and isotonic hand configurations, and 9 basic movements of the wrist). To collect the data, each subject was asked to maintain the hand gesture for 5 seconds, followed by resting for 3 seconds, and the experiment is repeated 6 times for each gesture. In addition, the acquisition setup included 12 Delsys Trigno electrodes (8 electrodes were wrapped around the radio-humeral joints, two around the biceps and triceps, and two around the flexor and extensor digitorum superficialis). They are designed to record hand kinematics, dynamics, and the corresponding muscular activity, and the sensors were linked to a laptop responsible for signal data acquisition sampled at 2kHz. Using such benchmarked database allows us to ignore various factors, including the experimental conditions which could otherwise affect the recorded results.

**Fig. 1:**
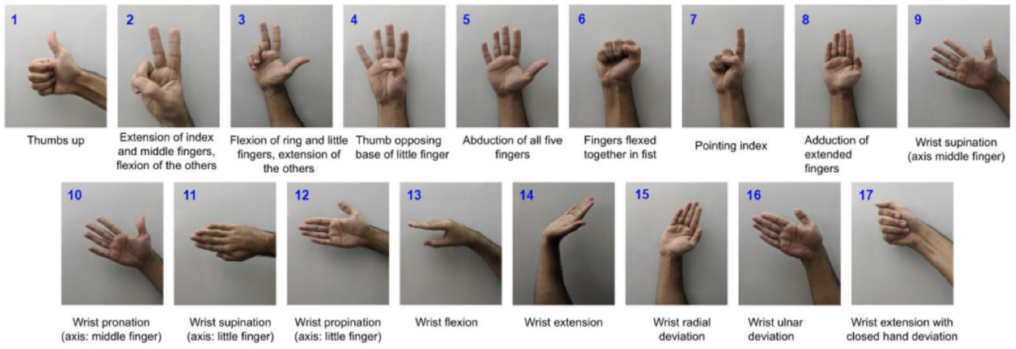
Total 17 gestures in DB2, Exercise Set B.

### B. Data Pre-processing

The three phases of repetition of voluntary muscles contraction form a trapezoidal profile and are visualized using sEMG accelerometer data. The upper base of the trapezoid is considered the steady-state of contraction, and the first slant of the trapezoid roughly outlines the transient phase. In this letter, we decide the length of the transient phase by the root mean square (RMS) of the accelerometer signals averaging across all subjects. As shown in Fig. 2, it is observed that the average RMS of the accelerometer data becomes steeper in the first 20% (1 second) of the gesture repetition. This part can be considered as the transient phase, and thus, we extract the first 20% of each repetition length to obtain the transient, discarding the remaining part of the repetition. In this way, the resulting data will have a total duration of 68 seconds (4 repetitions * 1 second per repetition * 17 gestures) for each subject.

**Fig. 2:**
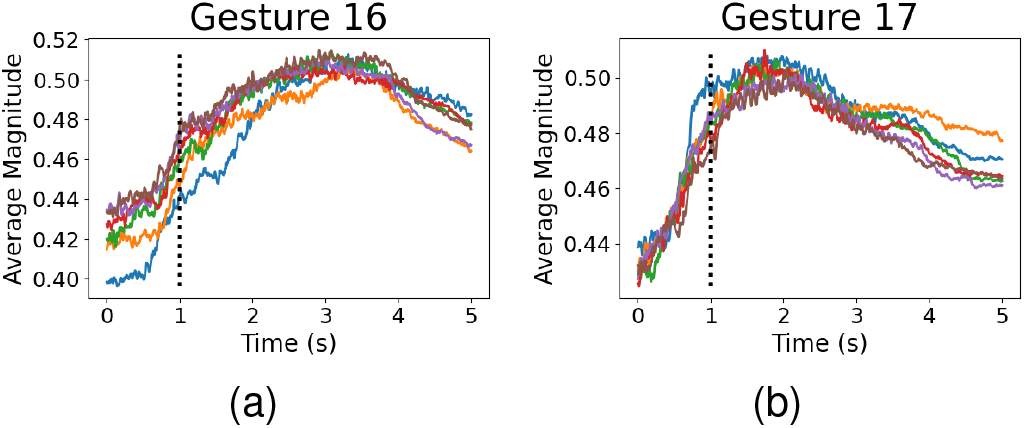
Illustration of transient phase signals: average root mean square of accelerometer data across all subjects for (a) gesture 16, and (b) gesture 17. The dotted line indicates that the transient phase ends at 1s. Repetitions are denoted by different colors.

A minimal preprocessing pipeline that includes normalization and rectification is then processed on the data. It normalizes the signals by using Z-score normalization with means and standard deviations from the training data and rectifies the normalized signals by taking the absolute values. Normalized and rectified signals are windowed with 300ms, and labels are assigned to each window. The signal data after windowing will be in the shape of 600*12. The 600 here represents the number of timesteps (300ms*2 kHZ=600), and 12 represents the number of sensors or channels. Based on the literature, 300ms is the largest window size required for realtime control. We use a window of size 300ms [25] with an overlap of 10ms to generate the training and testing sets. The data in DB2 dataset is already Hampel filtered to remove 50Hz powerline interference [26].We do not apply any extra lowpass filtering techniques to the signal since it reduces the quality of the signal. For train-test split, we leverage repetitions 1,3,4 and 6 for training the deep learning model and repetitions 2,5 for validating the trained model [26]. The transient phase of the signal being a small part of the data sample drastically reduces the time required to train the model and results in a lower calibration time.

## III. Model Architecture

We propose the concept of Dilated Efficient Capsule Network in this letter for sEMG signals classification, and apply transfer learning on it to improve gesture-prediction accuracy and minimize the needed data for training.

In the literature, Convolutional Neural networks (CNN) have helped achieve remarkable results in problems including image classification [27]–[29] to object detection [30]–[32]. However, translation invariance achieved by the CNN comes at the expense of losing some information of the object’s location. To counterbalance the problem, we propose our new model architecture involving Capsules which are vector representations of features. Moreover, each capsule involved in the network dynamically describes how the entity is instantiated. Hence, we drop the working principle of traditional neural networks where the scalar unit is activated based on its synergy with the learned feature detectors. Capsule networks also leverage the concept of Routing by Agreement, where the predictions of low-level capsules are routed to their best match parent. This helps assess the reciprocal agreement between groups of neurons to capture covariance and lead to a compact model with fewer parameters and better capability to generalize on new data.

The model architecture of the proposed Efficient CapsNet has two basic building blocks as described in Fig. 3: a) the Convolutional block, and b) the Capsule block. The Convolutional block holds four CNN layers with each having a 2D convolutional layer followed by a Dropout Layer and Batch Normalization Layer. The first two CNN blocks have 128 filters, and the other two have 256 filters. A Relu activation function is used after each convolutional layer. We use the Dropout regularization technique to avoid the possible case of over-fitting. The dropout rate in our model is set to 0.5. We have the Convolutional block followed by a Capsule block. The Capsule block involves a “Primary Conv” layer that has a 2D convolutional layer followed by a reshape function. It is necessary to ensure that the length of the vector lies in the range from 0 to 1 because it represents the probability of information routing from the current layer to the next capsule layer. Hence a “squash” activation is used which drives the length of the large vector towards 1 and the small vector towards 0. The dimension of each capsule and the number of channels is set to 4. The Primary Conv layer is followed by the Capsule layer, where the Routing algorithm works. This layer just expands the output of the neuron from a scalar to a vector. The number of routing iterations is assumed to be 3. The number of capsules in the final Capsule layer would be equivalent to the number of gestures being predicted, 17. The proposed model then outputs the probability for each of the 17 gestures.

**Fig. 3:**
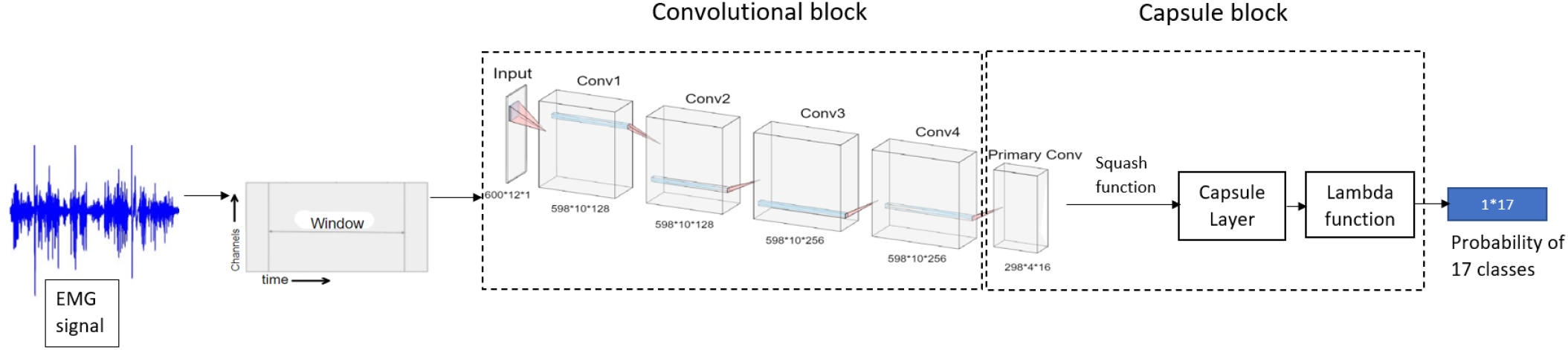
The Illustration of the model architecture of Dilated Efficient CapsNet. 1) We process the sEMG signals by windowing, and make it the input of Dilated Efficient CapsNet. 2) The Dilated Efficient CapsNet consists of two blocks: the convolutional block and the capsule block. The Convolutional block has four Conv layers, each having a 2D Convolutional Layer followed by a Dropout Layer and Batch Normalization Layer, with a Relu activation function at the end. The Capsule block has a Primary Conv layer that has a 2D Convolutional Layer followed by squash function, a Capsule layer to perform routing algorithms, and a Lambda Function to generate the output. 3) The output of this model architecture is the probability of each signal classified as one of the 17 gestures.

The Efficient CapsNet is an improvement of the currently existing CapsNet model in literature [33]. The original CapsNet had 16.3M trainable parameters in comparison to the Efficient CapsNet, which has 1.7M trainable parameters. The primary difference between the two models is their architecture. The original CapsNet has a single convolutional layer in the convolutional block in comparison to the Efficient CapsNet, which has four convolutional layers, each with a large number of filters. To avoid the complexity of the original CapsNet model, the Efficient implementation removes the reconstruction/decoding block of the CapsNet. This reduces the number of trainable parameters and training time significantly. Removing the decoder part does not affect the model’s performance because we currently use minimal preprocessed windowed data as the input to the model and not images. To improve the performance of the model, we make use of dilation to expand the area of reach without pooling. We use a dilation rate of (2,2) in the 2nd convolutional layer followed by (4,4) in the 3rd layer and (8,8) in the last layer of the convolutional block. The total number of trainable parameters for the Dilated Efficient CapsNet becomes 3.7M, however, the additional padding incorporated with dilation allows the center of the kernel to pass over the edge channels, providing additional useful information. The training process in a real-life situation can be quite time-consuming. To solve this problem, we introduce the concept of Transfer learning and apply it in our research to try to achieve a higher classification accuracy with comparatively fewer data. Since the effectiveness of the transfer learning hinges on the quality of the subset of the data used to initialize the majority of our model layers, a poor subset would not accurately represent the inter-subject variance and not be very useful. We list the top 5 performing subjects among all the 40 subjects by the individual scores obtained after training the model per subject. The Dilated Efficient CapsNet model is then trained on the data of the top 5 subjects before it is completely frozen, excluding the first two layers of the convolutional block. Leaving the first few layers trainable, allows our model to learn the fine features of unseen subject data. The new model can learn from the common knowledge of best-performing subjects and then continuously learn from new inputs when there is fresh information coming in. In this way, we can reduce the negative impact of less training data, improve training efficiency, and even achieve better classification accuracy.

## IV. Experiments and Results

A set of experiments are conducted to validate our proposed model architecture and the segmentation scheme of the transient phase. We compare the performance of Dilated Efficient CapsNet (Dilated Eff-Caps) and Transfer Learning Dilated Efficient CapsNet (Top5 Eff-Caps) with MLP, 2D CNN, and RNN-CNN Hybrid [34] by predictive accuracy, and compare all the models by different transient segmentation scheme with different data volume.

The Hybrid model architecture was inspired by [34]. The hybrid model has three components: the LSTM block composed of four LSTM layers, each with 128 units; the CNN block with seven convolutional layers with the filters of 32, 64, 64, 128, 128, 256, respectively and dilation rate on CNN layers two through six of 2, 2, 4, 8, 8, 8; and the classifier block, three fully connected dense layers with units 64, 32, 17. The size of the MLP and 2D CNN network was created to be comparable to the number of trainable parameters in the Hybrid model 1.7M. The MLP architecture used is a simple five layer network with the following number of units per layer: 256, 128, 64, 32, 17. The 2D CNN was composed of four 2D convolutional layers with filters of 32, 32, 64, 64. The convolutional block was then followed by a small classifier block of two dense layers with 32 and 17 units.

Regarding data segmentation scheme, we have temporal segmentation and repetition segmentation. As shown in Fig. 4, we split sEMG signals by repetition 1,3,4,6 and repetition 2,5. Purple repetitions 1,3,4,6 are fed into the models as training samples, and red repetitions 2,5 are served as the testing data. To validate the proposed model’s ability to achieve high predictive accuracy with much less training data, we extract different percentages of transient signals as model input. Temporal segmentation extracts signals via horizontal chronological order, while repetition segmentation sees each repetition as 25% of the training transient signal and extracts a various number of repetitions based on experimental need via vertical order.

**Fig. 4:**
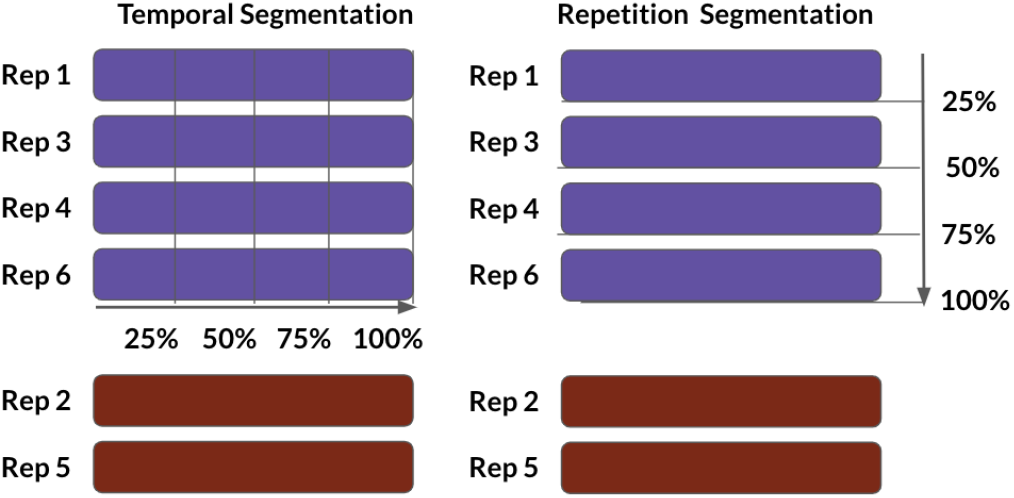
The illustration of data segmentation scheme: temporal segmentation and repetition segmentation. Repetitions in purple are training samples, and repetitions in red are testing samples. We extract different percentage of the transient phase via the direction of arrow.

**Fig. 5:**
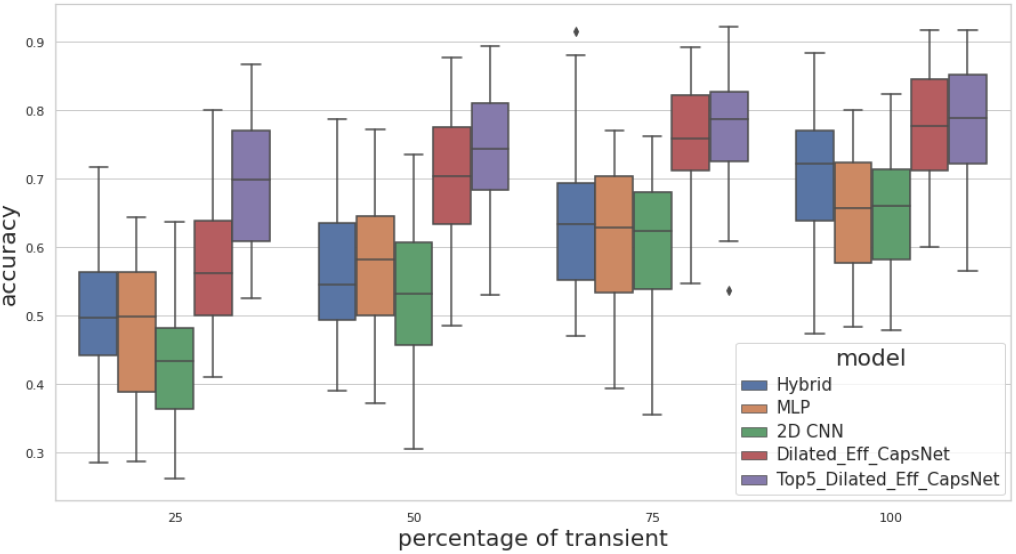
Temporal segmentation performance comparison across selected models

**Fig. 6:**
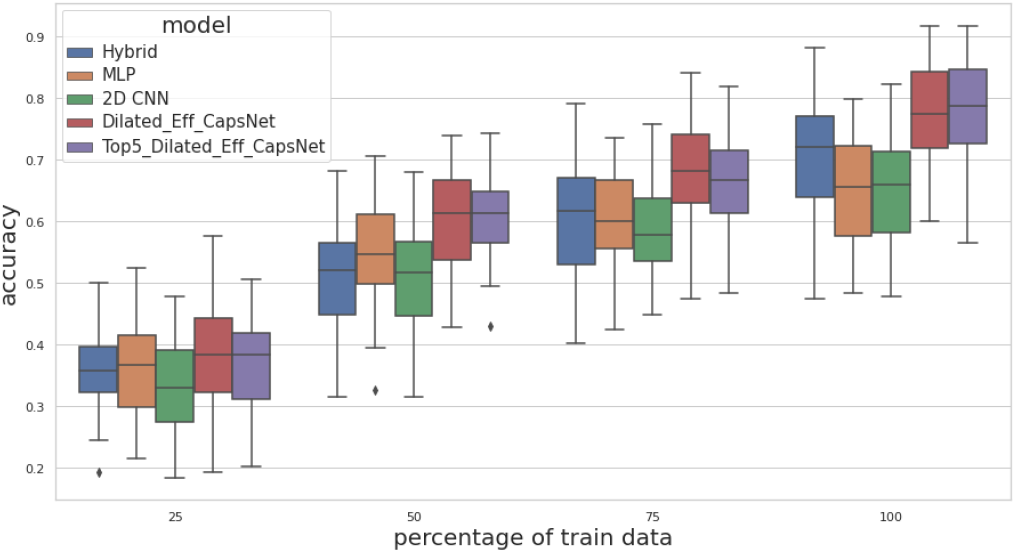
Repetition segmentation performance comparison across selected models

The comparative average model performance of temporal segmentation is included in the table 1, and repetition segmentation model comparison is included in the table 2. Both tables are followed with a box plot to further illustrate the results.

**TABLE I.** Results of Temporal Segmentation (average acc.)

| Transient Percentage | MLP | 2D CNN | Hybrid | Dilated Eff-Caps | Top5 Eff-Caps |
| --- | --- | --- | --- | --- | --- |
| 25% | 0.484 | 0.429 | 0.502 | 0.565 | 0.694 |
| 50% | 0.567 | 0.529 | 0.562 | 0.701 | 0.740 |
| 75% | 0.616 | 0.603 | 0.639 | 0.761 | 0.777 |
| 100% | 0.647 | 0.651 | 0.713 | 0.775 | 0.783 |

**TABLE II.** Results of Repetition Segmentation (average acc.)

| Transient Percentage | MLP | 2D CNN | Hybrid | Dilated Eff-Caps | Top5 Eff-Caps |
| --- | --- | --- | --- | --- | --- |
| 25% | 0.360 | 0.334 | 0.359 | 0.381 | 0.367 |
| 50% | 0.543 | 0.511 | 0.513 | 0.603 | 0.605 |
| 75% | 0.601 | 0.584 | 0.606 | 0.680 | 0.663 |
| 100% | 0.647 | 0.651 | 0.713 | 0.775 | 0.783 |

Overall, we can notice a higher predictive accuracy of temporal segmentation experiments than repetition segmentation experiments. All models’ predictive accuracy of repetition segmentation tasks are highly correlated to the amount of data fed as training samples, while in temporal segmentation, Dilated Eff-Caps and Top5 Eff-Caps can still achieve high accuracy with much less data.

Based on temporal segmentation, Top5 Eff-Caps can achieve 69.4% predictive accuracy with only 25% of the transient (5% of the complete signal), and achieve 74%, 77.7%, 78.3% with 50%, 75%, and 100% percentage data respectively. The Top5 Eff-Caps outperforms any other model architecture in terms of the predictive accuracy with any transient percentage. At most, it can achieve 43.4% higher accuracy with 25% of transient data compared to MLP; at least, it can achieve 10% higher accuracy with 75% of transient data compared to the Hybrid model.

Based on repetition segmentation, the effectiveness of transfer learning is not as dominant as before. Some of the Top5 Eff-Caps have lower accuracy compared to simple Dilated Eff-Caps. However, in terms of Dilated Efficient CapsNet Architecture, both Dilated Eff-Caps and Top5 Eff-Caps still outperform all other comparative models, validating the effectiveness of the proposed model architecture in this letter.

## V. Conclusion

This letter proposes transfer learning of Dilated Efficient CapsNet to optimize the gesture prediction accuracy with much less data provided compared to conventional studies. Instead of using complete sEMG signal repetitions, we utilize only the transient phase as the training samples. In this way, we reduces the training data to at least 20% of the original and transform the traditional classification problem into prediction. Utilizing only transient signals brings the research a step closer towards real-time predictive applications in large-scale neurorobotics. In addition, with the proposed Dilated Capsule Neural Network being introduced to this field, its inherent characteristics of saving features as vectors help us capture the correlated information within signals and across repetitions. It ensures the performance of proposed model architecture with very few data compared to conventional deep learning methods. Via experiments, Transfer Learning based Dilated Efficient CapsNet can achieve an average predictive accuracy of 78.3% across 35 subjects on transient signals (20% of the complete signal repetition). Even with 25% of the transient (5% of the complete signal repetition), it can still achieve an accuracy of about 70%.

In this work, we also examine two different data segmentation schemes by comparing the performance of various models under these two circumstances: temporal segmentation and repetition segmentation. By processing with these two segmentation schemes, we can check the feasibility of further reducing the training data by 25%, 50%, and 75% of its original. A superiority of temporal segmentation over repetition segmentation is observed, and the effectiveness of transfer learning under temporal segmentation is larger than that under repetition segmentation. Though we use much fewer data than previous research, it is noteworthy to mention that in our study, all data had been collected from able-bodied subjects. The neurophysiology of a healthy population can hardly reflect amputees’ bio-electrical conditions, and various amputations would also lead to greater challenges of training only on transient signals of non-intact subjects. Hence, we will collect the sEMG data from amputees in the future and validate our model architecture in more practical applications.

